## Supplementary figures and images for "Sclerostin Small Molecule Inhibitors Promote Osteogenesis by Activating Canonical Wnt and BMP Pathways"

### Fig 1 - figure supplement 1

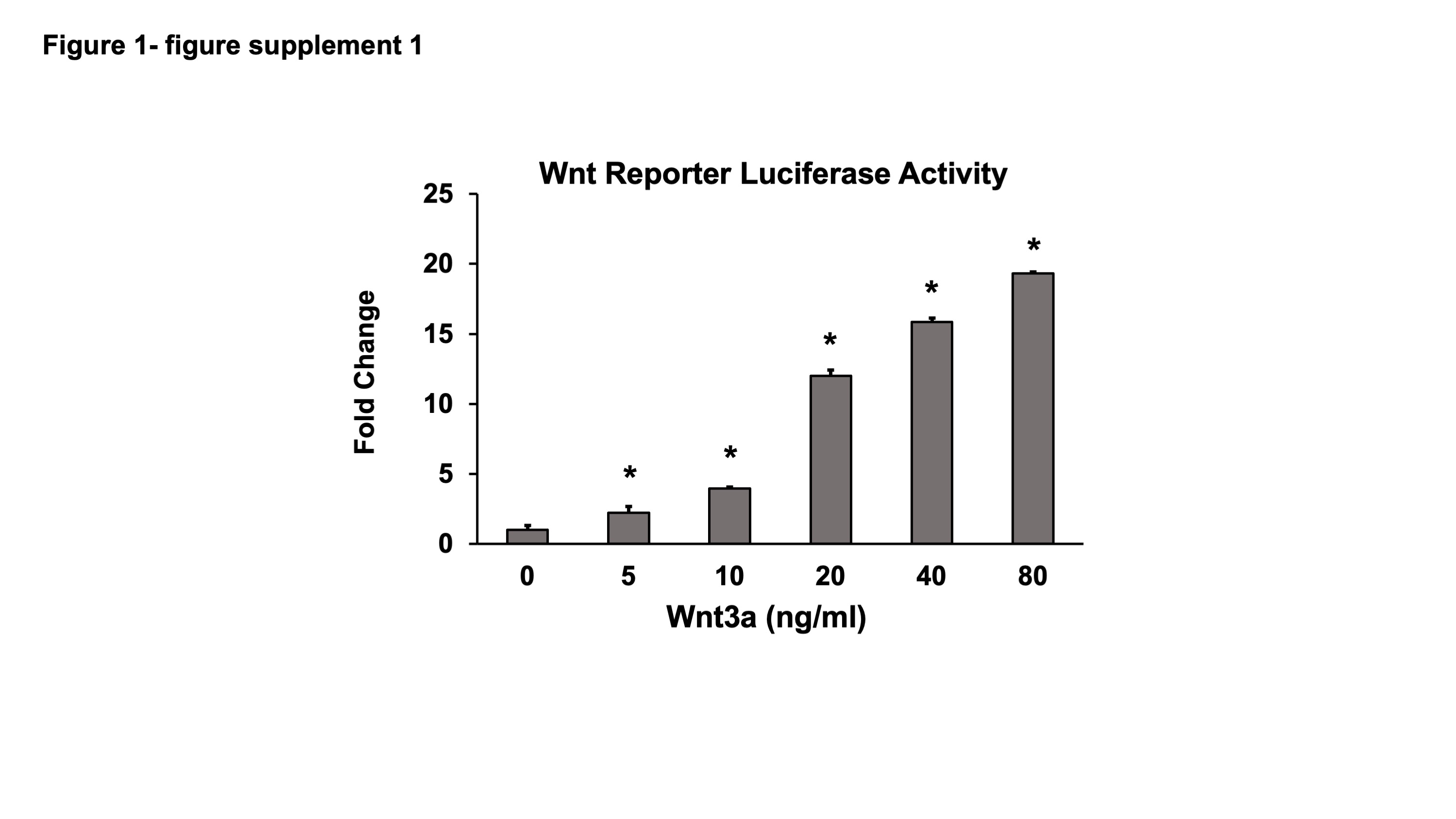

### Fig 2 - figure supplement 1

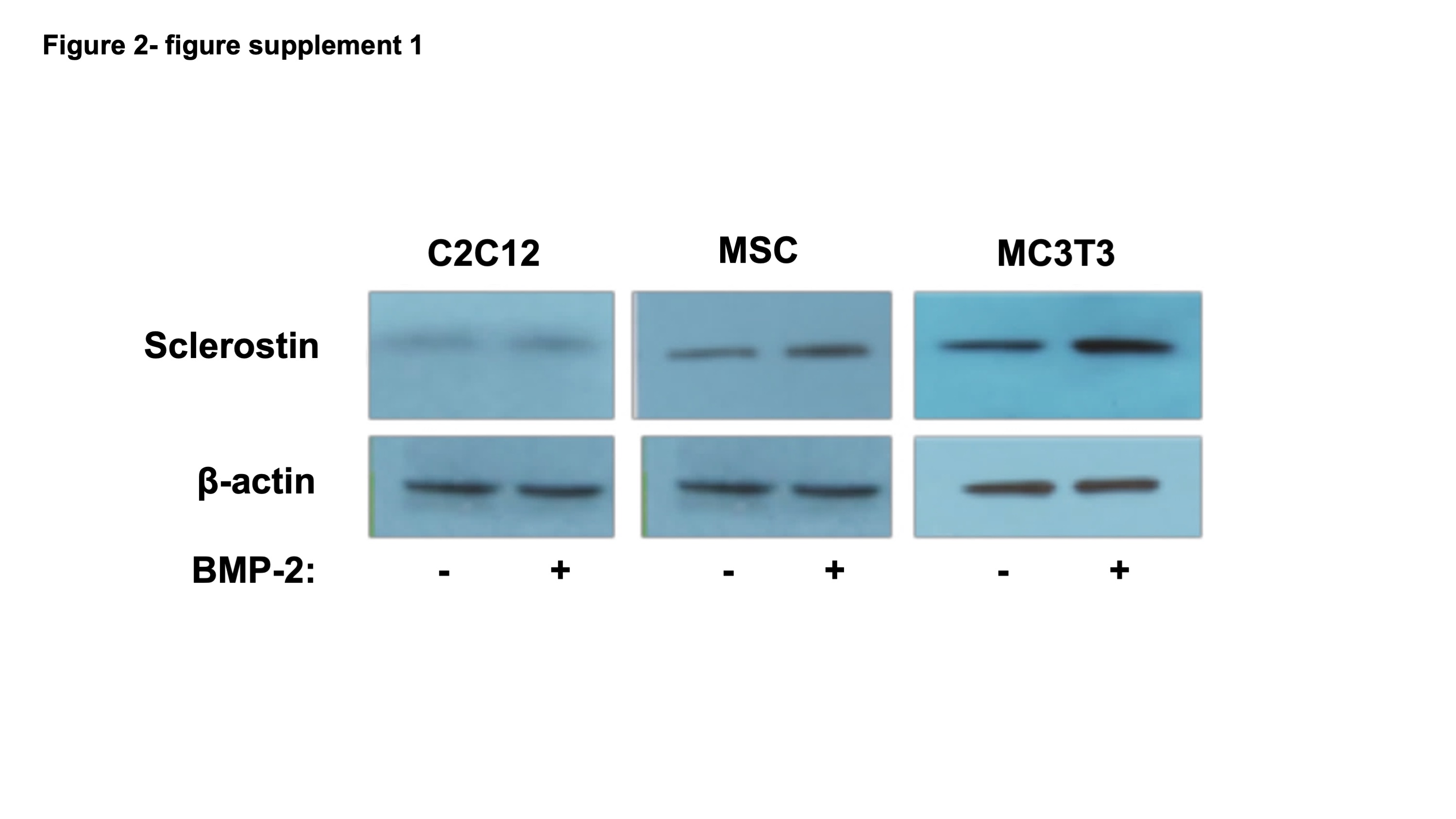

### Fig 9 - figure supplement 1

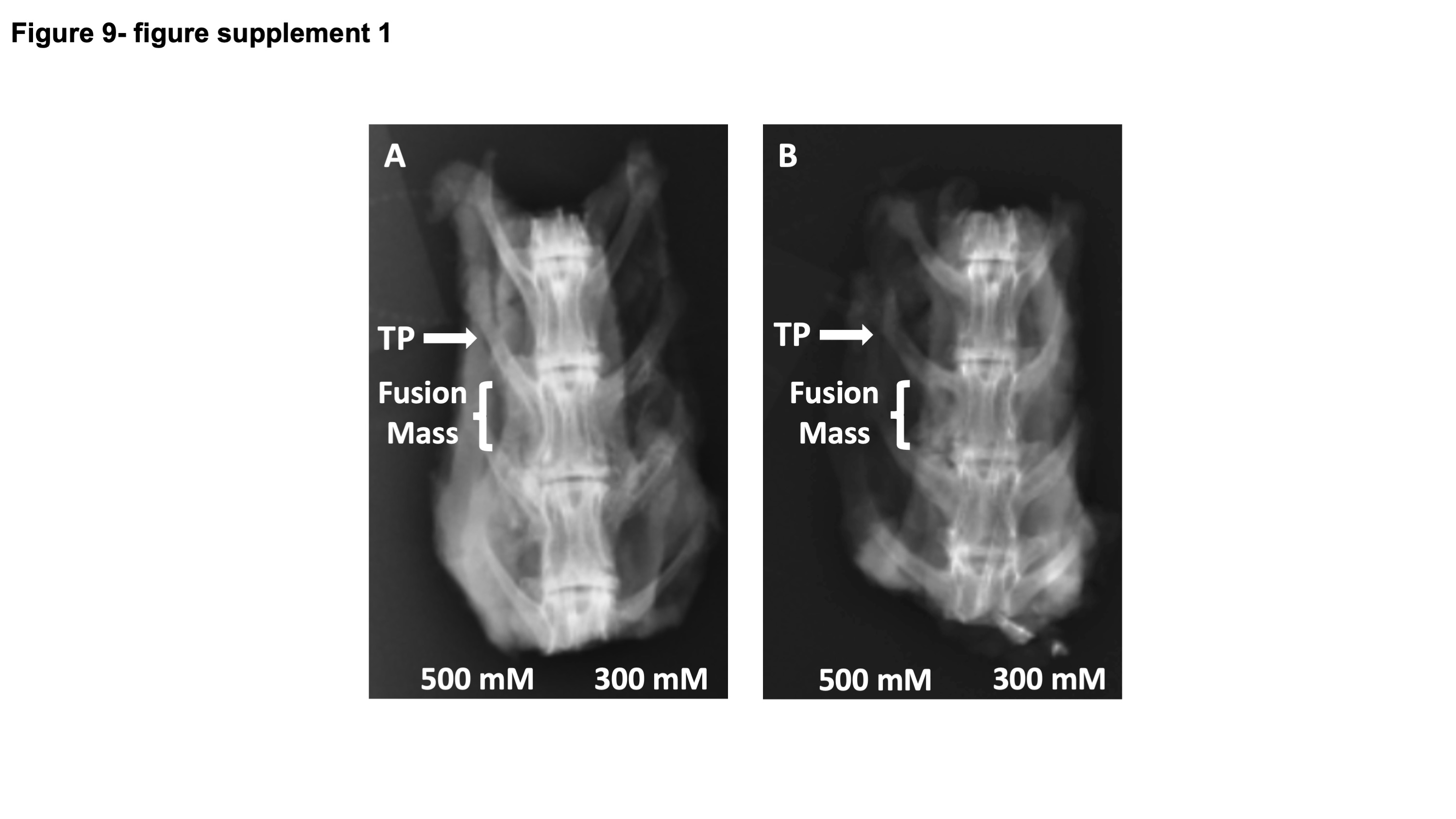
